# Lack of Ameliorative Effect of Chronic Oral Nicotine on Olfactory Dysfunction in Young Adult Presenilin-1/2 Double Knockout Mice of a Dementia Model

**DOI:** 10.1101/2024.11.14.623574

**Authors:** Youwen Si, Shuai Qiu, Jingyan Jia, Jiale Duan, Wenbo Feng, Bo Meng, Feiyan Qi

## Abstract

Olfactory dysfunction is a prevalent and early feature of neurodegenerative diseases including Alzheimer’s disease while the features and underlying mechanisms remain inadequately understood. This investigation aimed to elucidate the olfactory function characteristics of presenilin-1/2 double knockout (DKO) mice, an established mouse model of dementia. Through a series of behavioral tests on DKO and wildtype (WT) mice at 2 and 6 months of age, we assessed their abilities in odor detection, discrimination, and olfactory memory. The findings revealed a substantial deficit in odor detection in 2-month-old DKO mice compared to WT mice, with a noteworthy age-related deterioration observed from 2 to 6 months. Although DKO mice exhibited normal odor discrimination and olfactory memory capabilities at 2 months, these functions declined with age, falling significantly lower levels than observed in WT mice by the age of 6 months. Subsequently, 2-month-old DKO and WT mice underwent a 4-month chronic nicotine treatment through drinking water. This intervention failed to ameliorate olfactory dysfunctions in DKO mice and exhibited no significant impact on WT mice, despite a discernible trend of a potential negative effect. In summary, this behavioral study elucidates that Presenilin-1/2 double knockout impairs odor detection in young adult mice and accelerates declines of olfactory discrimination and memory. The chronic administration of nicotine did not effectively mitigate olfactory function deficiencies in young adult DKO mice, and the results also highlight age as an important contributing factor.

## 1. Introduction

Dementia is a multifaceted neurological disorder impacting memory, cognitive processes, and behavior. According to the World Health Organization (WHO) in 2023, over 55 million individuals worldwide are affected by dementia, with nearly 10 million new cases reported annually. Projections estimate that by 2050, the global prevalence of dementia will rise to 152 million. Olfactory function, or the sense of smell, plays a crucial role in connecting animals, including humans, to their environment and is integral to various activities of daily living such as odor detection, recognition, memory, and discrimination. Odor detection involves the nose’s ability to identify different scents, olfactory recognition pertains to identifying specific smells, and olfactory memory involves the capacity to remember and recall odors from past experiences. Olfactory recognition, a subset of olfactory memory, entails identifying previously encountered odors. Olfactory discrimination is the ability to differentiate between different odors. Overall, olfactory function is an important sense enabling animals to detect, identify, and recall odors. However, olfactory dysfunction is common in neurological disorders, for example, Alzheimer’s disease (AD), the most usual form of dementia which may contribute to 60-70% of cases. A reduced smelling ability (known as hyposmia), odor identification and discrimination dysfunction have been observed in AD and mild cognitive impairment (MCI) patients ^1^. Furthermore, the olfactory system is often one of the earliest to show dysfunction and thought to be a predictor of disease progression ^2,3^. It is noteworthy that more than 15 million people worldwide experience persistent COVID-19-related olfactory dysfunction, possibly caused by viral invasion of olfactory bulb, and have been linked to initiate a cascade of neurodegeneration akin to AD and Lewy body disease ^4^.

While the causes of olfactory dysfunction in neurodegenerative diseases remain unclear, factors such as inflammation and oxidative stress are suspected. Amyloid β (Aβ) deposition and Tau tangles, two hallmarks of AD pathology, are associated with olfactory deficiency. While Aβ deposition have been detected in the olfactory bulb of AD patients ^5^, indicating a potential role in olfactory impairment, using Pittsburgh compound B-positron emission tomography (PiB-PET), have not found a direct correlation between Aβ burden and olfactory dysfunction ^6^. Similarly, Tau hyperphosphorylation and neurofibrillary tangle formation have been observed in the olfactory bulb of AD or PD patients ^5^; in P301S tau transgenic mice, olfactory sensitivity was significantly impaired at 3 months of age with noticeable progressive tauopathy in the olfactory bulb and piriform cortex ^7^. In contrast, no olfactory impairment was detected in THY-Tau22 mice until 12 months although Tau pathology early affected olfactory cortical structures ^8^. The discrepant evidence suggests an ambiguous causal relationship and the complexity of underlying mechanisms.

Presenilin-1(PS1) and presenilin-2 (PS2) are important components of γ-secretase which is responsible for cleaving transmembrane proteins, including amyloid precursor protein (APP) and Notch receptors ^9^. Furthermore, PS1 and PS2 play roles in γ-secretase-independent functions, such as stabilizing β-catenin in Wnt signaling pathway, regulating calcium homeostasis, interacting with synaptic transmission, and regulating the autophagy-lysosome system ^10^. Studies have shown that mutations in the *APP* and/or *PS1*/*PS2* gene are the main causes of early-onset AD which leads to increasing Aβ deposition and further amyloidogenic pathology ^11^. Interestingly, Feng et al. crossed the forebrain-specifically *PS1* knocked out mice with systemically *PS2* gene knocked out mice and obtained the conditional *PS1*/*PS2* double gene knockout (DKO) mice ^12^, which showed a series of neuro-degenerative symptoms including specific forebrain atrophy with age, tau protein hyperphosphorylation, massive neuronal apoptosis, glial cell proliferation and cognitive dysfunction, but without Aβ deposition as the absence of the functional γ-secretase. The DKO mouse is a distinctive non-amyloidogenic model for dementia, providing valuable insights into the exploration of non-beta-amyloid related function of Presenilins. Although our previous studies have exhibited the cognitive, non-cognitive and emotional symptoms in DKO mice ^13-15^, the potential aberrations in the olfactory function of this mouse model remain unexplored.

The olfactory bulb, modulated by the central cholinergic system, plays a key role in olfactory processing^16^. Studies have shown that forebrain cholinergic denervation is closely related to olfactory deficits in neurodegenerative diseases, especially in Alzheimer’s disease (AD)^17^. It is notable that the medicinal effects of nicotine, the main components of tobacco, on neuro-degeneration diseases treatment are obscure as the olfactory decline might be associated with other toxic components in cigarettes exposures. Even the relationship between nicotine itself and neurodegenerative diseases is debatable. Nicotine acts as an agonist of nAChRs which can further regulate the release of a bunch of neurotransmitters, such as dopamine, glutamate, γ-aminobutyric acid, 5-HT, norepinephrine, therefore affects not only learning and memory but also addiction and mood ^18^. It has been well known that AD is also characterized by acetylcholine (ACh) system deficiency, including reduction of nicotinic acetylcholine receptors (nAChRs), attenuated activity of cholinergic synthetic or inactivating acetylcholinesterase ^19^. It is widely acknowledged that prolonged exposure to nicotine induces desensitization of nAChRs, resulting in cognitive deficits among otherwise healthy individuals ^20^. On the other hand, it was reported that transdermal nicotine administered to nonsmoking MCI patients over 6 months improved cognitive measures of attention, memory, and mental processing, but not in ratings of clinician-rated global impression ^21^. Moreover, nicotine has been demonstrated to modulate various signaling pathways beyond the cholinergic pathways, thereby introducing a higher degree of complexity to its functionality, potentially eliciting distinct effects in diverse situations ^22^.

In this study, we conducted behavioral assays to assess olfactory function in various age groups of DKO mice, examining the potential link between dementia and olfactory performance. Additionally, we evaluated the impact of chronic nicotine exposure through drinking water on young adult DKO and age-matched wildtype (WT) mice, aiming to build a theoretical foundation for early olfactory assessment as a potential diagnostic tool for dementia.

## 2. Methods and Materials

### 2.1 Animals and nicotine administration

DKO mice were generated by crossing forebrain PS1 knockout mice (CaMKII-Cre+, *PS1* flox/flox) with conventional *PS2* knockout mice (*PS2*-/-), as previously described ^12^. The littermate WT mice (CaMKII-Cre-, *PS1*-/-, *PS1*+/+) served as control mice, with both DKO and WT mice maintained on the C57BL6/J background. Mice were accommodated in cages with a population density ranging from 3 to 5 animals per enclosure, kept at 24 ℃ and 40-70% humidity with access to food and water *ad libitum* on a 12-h light/dark cycle (light on from 7:00 a.m. to 7:00 p.m.) and subjected to cage changes on a weekly basis in ECNU Public Platform for Innovation (010). Both male and female mice were used with approximately equal ratio. Behavior tests involved 2-month-old and 6-month-old WT and DKO mice, with sample sizes of n = 22∼36 for each group (see supplemental Table S1). For chronic nicotine administration, (-)-nicotine (Alta Scientific, China) was supplied to 2-month-old WT and DKO mice in the drinking water *ad libitum* at the dose of 100 μg/mL for 4 months (WT-N and DKO-N, respectively. n = 8∼10) and the age-matched WT and DKO mice supplied with drinking water served as blank control (WT-B and DKO-B, respectively, n = 10∼12). The Animal Ethics Committee of East China Normal University authorized all experiments.

### 2.2 Martials

2-Methylbutanoic Acid (2-MB) (98% purity, Cat. # M02949, Meryer Biotech, China), ginger pure essential oil (GTIN: 675235000219, Amphora Aromatics Ltd, UK), and cinnamon essential oil (ZRZR Official Flagship Store, China) were purchased for use in olfactory function tests (IACUC approval ID #M10020).

### 2.3 Behavior tests

#### Buried Food Test

Buried food test is one of the most classic olfactory behavior experiments, used to assess the ability of animals to detect odor ^23^. The experimental apparatus was a 35 cm × 20 cm× 15cm (L× W× H) cage with a 5 cm layer of bedding material. A standard food pallet was buried 1.5 cm below the surface of the bedding material in a random location. Each mouse was placed in the apparatus for 5 min daily to acclimate for 3 consecutive days. Mice were fasted for 12 hours before the test. During the experiment, each mouse was placed into the cage and the latency (in seconds) required for the animal to successfully retrieve the pellet was recorded, with a maximum time of 5 min. Failures were recorded as 600 seconds. Experiment was conducted for 3 consecutive days, with a new buried location each day, and the average latency was calculated.

#### Olfactory Avoidance Test

The olfactory avoidance test measures the avoidance time of mice against the disgusting odor 2-methylbutyric acid (2-MB) (2-MB, a pungent odorant of spoiled foods, makes mice disgust but does not produce fear) and detects the ability of mice to distinguish odors ^24^. The apparatus was a 35 cm × 20 cm× 15cm (L× W× H) cage without bedding material. An opaque partition divided the cage into two compartments, with the larger compartment having double the volume of the smaller one. There was a hole (5 cm × 5 cm) at the bottom of the partition, allowing mice to move freely. In the acclimatization stage, each mouse spent 5 minutes in the apparatus with a filter paper (2 cm × 2 cm) moistened in the small compartment for 2 consecutive days. In the training stage, each mouse spent 5 minutes in the apparatus with a filter paper moistened with 20 μL distilled water, and the time that the mouse stayed in the large compartments within 3 minutes was recorded as T1; then in the test stage, the filter paper was replaced with a filter paper soaked in 20 μL2-MB, and the time that the mouse stayed in the large compartments within 3 minutes was recorded as T2. After testing each mouse, the cage was thoroughly cleaned with 10% ethanol and then rinsed meticulously with clean water to eliminate any residual and ensured complete drying of the testing arena by employing clean towels. Avoidance time was calculated as T2-T1.

#### Olfactory Memory Test

The olfactory memory experiment is modified from the traditional three-box experiment, which can comprehensively investigate the ability of mice to distinguish and remember odors^25^. The apparatus was a 55 cm × 30 cm× 25 cm (L× W× H) cage with three compartments of equal size, each with a 5 cm×5 cm hole for movement. In both left and right compartments there was a petri dish with bedding material, alongside a 2 cm×2 cm filter paper dipped in either 10 μL cinnamon essential oil or 10 μL ginger essential oil. According to our preliminary test, mice showed no obvious preference for cinnamon essential oil or ginger essential oil. Mice were fasted for 12 h before the experiment. The first 3 days of the experiment are the training phase. A piece of food pallet was buried in the petri dish on the same side with ginger essential oil filter paper. The mouse was put into the middle compartment and allowed to move freely. After 5 minutes, the mouse was removed, and the cage was cleaned for the next mouse. On the fourth day, the petri dish was not buried with food. After the mouse was placed in the middle compartment, the time it stayed in each of the three compartments within 5 minutes was recorded. We calculated the preference index by assessing the time spent with the cued odor relative to the total time, where a 50% score indicates neutrality.

#### Data analysis and Statistics

Behavior tests were monitored and recorded by a camera, and the Anymaze software (Stoelting, U.S) was used for data analysis. Data were expressed as Mean ± S.E.M., with statistical significance calculated using Student’s *t*-test, Two-way ANOVA or repeated measure ANOVA in SPSS software. Statistical significance levels were set as follows: *P < 0.05, **P < 0.01, ***P < 0.001.

## 3. Results and discussion

### 3.1 Age-Dependent Decline in Olfactory Function in DKO Mice

DKO mice have been reported to exhibit various age-dependent neuron-degenerative phenotypes ^13^. At 2 months old, DKO mice showed relatively normal recognitive capability and normal noncognitive species-typical and home-cage behaviors. However, at 6 months old, DKO mice showed significant cognitive and non-cognitive capabilities deficiency as well as emotional disorder. In this study, we performed a serial of olfactory function tests on 2-month-old and 6-month-old DKO mice, with the comparison with the age-matched WT mice.

The Buried food test was applied to measure the odor detection capability. The overall statistical analysis reveals a significant genotype effect *F* _(1,115)_ = 80.035, P < 0.001; the age effect *F* _(1,115)_ = 36.197, P < 0.001 and genotype × age effect *F*_(1,115)_ = 22.021, P < 0.001. Specifically, at 2 months old, WT mice spent 83.1 ± 4.2 sec in digging out the food pallet, significantly less time than DKO mice, which spent 117.2 ± 6.3 sec (P < 0.05); at 6 months old, WT mice spent 93.8 ± 4.1 sec, while DKO mice spent a significantly longer 204.1 ± 16.1 sec (P < 0.05) (Figure 1A), which indicates that both 2-month-old and 6-month-old DKO mice showed significant longer latency of digging out the food pallet than that in WT mice; Further stratified analysis based on genotype reveals that the age effect is significant for DKO mice (P < 0.001), which means the age-dependent odor detection capability decline in DKO mice, whereas no significant difference between the latency in 2-month-old and 6-month-old WT mice (P > 0.05). The buried food test suggests DKO mice exhibit a progressive decline in odor detection, which worse with age.

**Figure 1.**
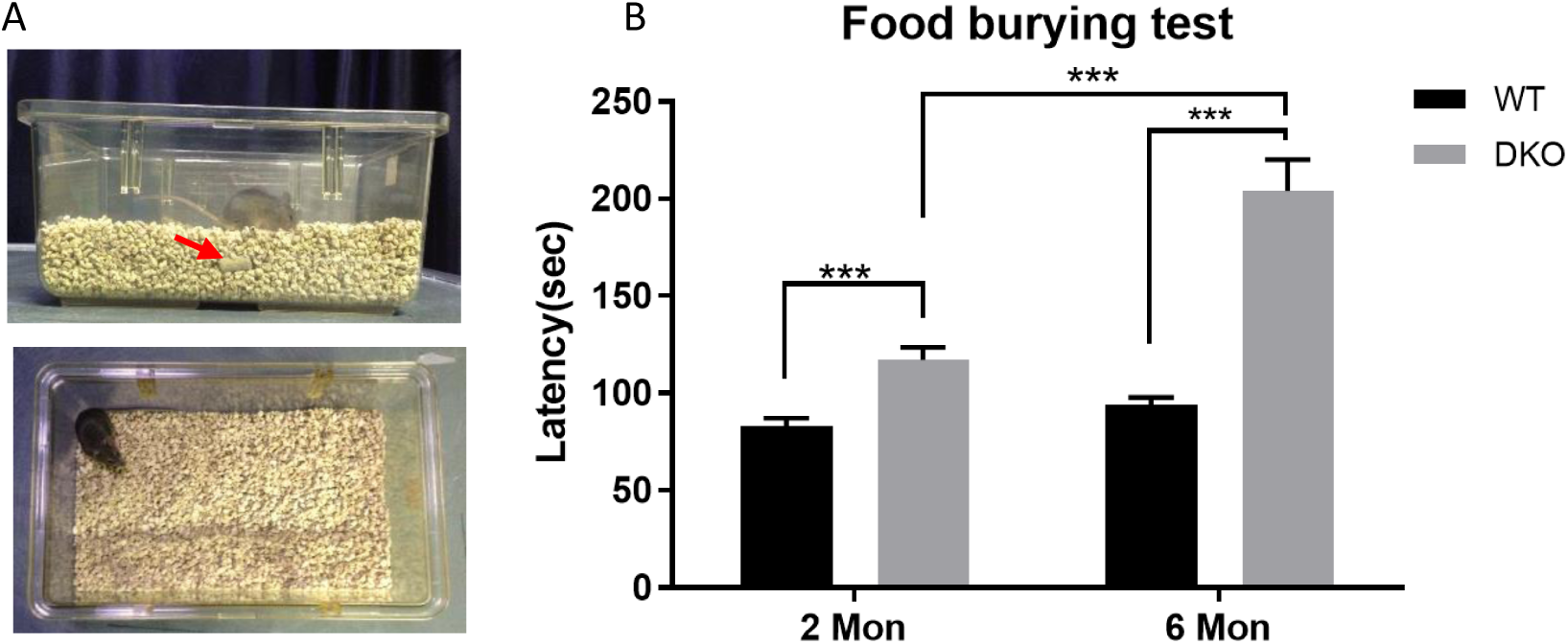
Buried food test. A. The apparatus used in the experiment. The red arrow showed the food pallet buried about 1.5 cm under the bedding material. B. The latency of the 2-month-old (2 Mon) and 6-month-old (6 Mon) WT and DKO mice. ***, P < 0.001.

The odor recognition capability was measured by the olfactory avoidance test. When the large compartment was exposed with disgusting odor 2-MB, the mice with normal odor recognition capability will spend less time in this area compared to when the compartment with neutral odor (water). Statistical analysis showed a significant genotype effect *F* _(1,115)_ = 10.029, P = 0.002, but no age effect (*F* _(1,115)_ = 2.190, P =0.142). There was, however,a genotype × age effect (*F* _(1,115)_ = 11.184, P <0.001), indicating a primary effect of genotype. At 2 months old, there was no significant difference in avoidance time between the DKO and WT mice (30.2 ± 6.4 sec and 28.7 ± 8.0 sec, respectively, P > 0.05), indicating normal odor recognition capability in DKO mice at 2-month-old. However, the 6-month-old DKO mice spent significantly less time (-4.9 ± 6.3 sec) in the large compartment than WT mice did (43.1 ± 9.3 sec, P < 0.001), suggesting a significantly loss of odor recognition in DKO mice. The olfactory avoidance test suggests that DKO mice experienced a decline in age-dependent odor recognition capabilities, exhibiting a significant deficiency in distinguishing odors compared to WT mice at 6 months of age. Similar to the buried food test results, WT mice did not demonstrate a noticeable age-dependent change in odor recognition capabilities within the first 6 months of age (P > 0.05). (Figure 2)

**Figure 2.**
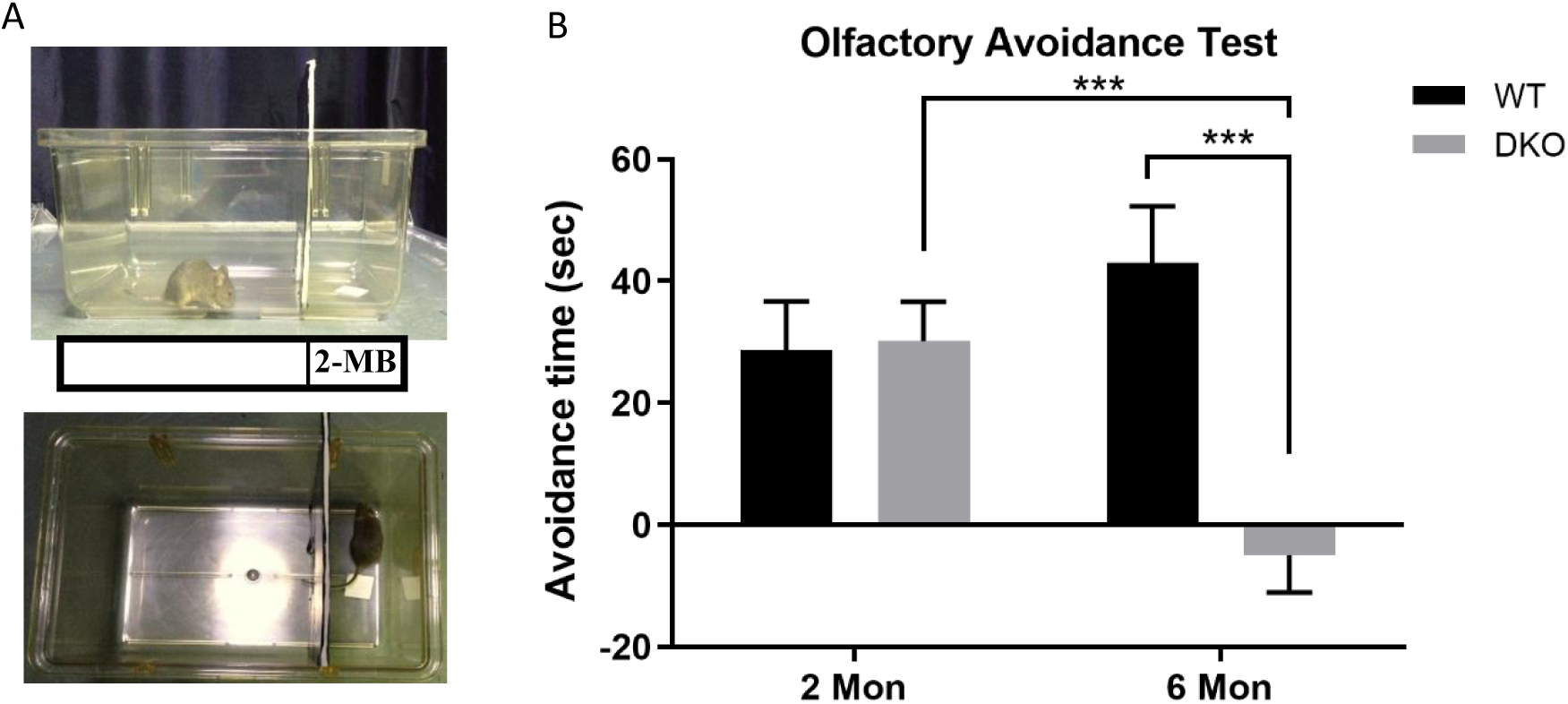
Olfactory avoidance test in 2-month-old and 6-month-old WT and DKO mice. A. The apparatus. The filter paper dipped in 2-MB was placed in the test stage. B. Avoidance time results. ***, P < 0.001.

Lastly, the olfactory memory test was conducted. In this test, the right compartment, associated with ginger odor, was linked with the presence of food during training. Mice with intact olfactory memory would exhibit a preference for this compartment during the test phase. Statistical analysis revealed a significant genotype effect *F* _(1,115)_ = 5.488, P = 0.021, but no age effect (*F* _(1,115)_ = 2.193, P = 0.141) or genotype × age effect (*F* _(1,115)_ = 1.357, P = 0.246). In detail, at the age of 2 months, both WT and DKO mice showed higher preference index (56.6 ± 2.1% and 57.3 ± 3.3%, respectively) and there was no significant difference (P > 0.05), indicating comparable olfactory memory; at 6-month-old, the preference index of WT mice remained consistent with 2-month-old WT mice (54.0 ± 1.7%, P > 0.05), while the preference index of DKO significantly declined (47.3 ± 1.1%, P < 0.05). There was also a significant difference between the two genotypes (P < 0.01), indicating an age-related decline in olfactory memory of DKO mice.

Across all three tests, neither 2-month-old nor 6-month-old WT or DKO mice showed any gender differences in performance (See supplemental figure S1, S2 and S3).

### 3.2 Chronic oral nicotine administration failed to rescue the olfactory deficiency in young adult DKO mice

Chronic nicotine administration was provided via drinking water at the dose of 100 μg/mL to 2-month-old WT and DKO for 4 months. Weekly body weight measurements showed no significant effect of nicotine on the body weight of either WT or DKO mice (P > 0.05) (Figure 4A). Drinking consumption was also recorded for 12 weeks (the measurement stopped when the behavior tests began) to exclude the possible impact on water intake. No significant differences were found among the four groups in drinking consumption (P > 0.05, Figure 4B), confirming equal nicotine intake across groups.

**Figure 3.**
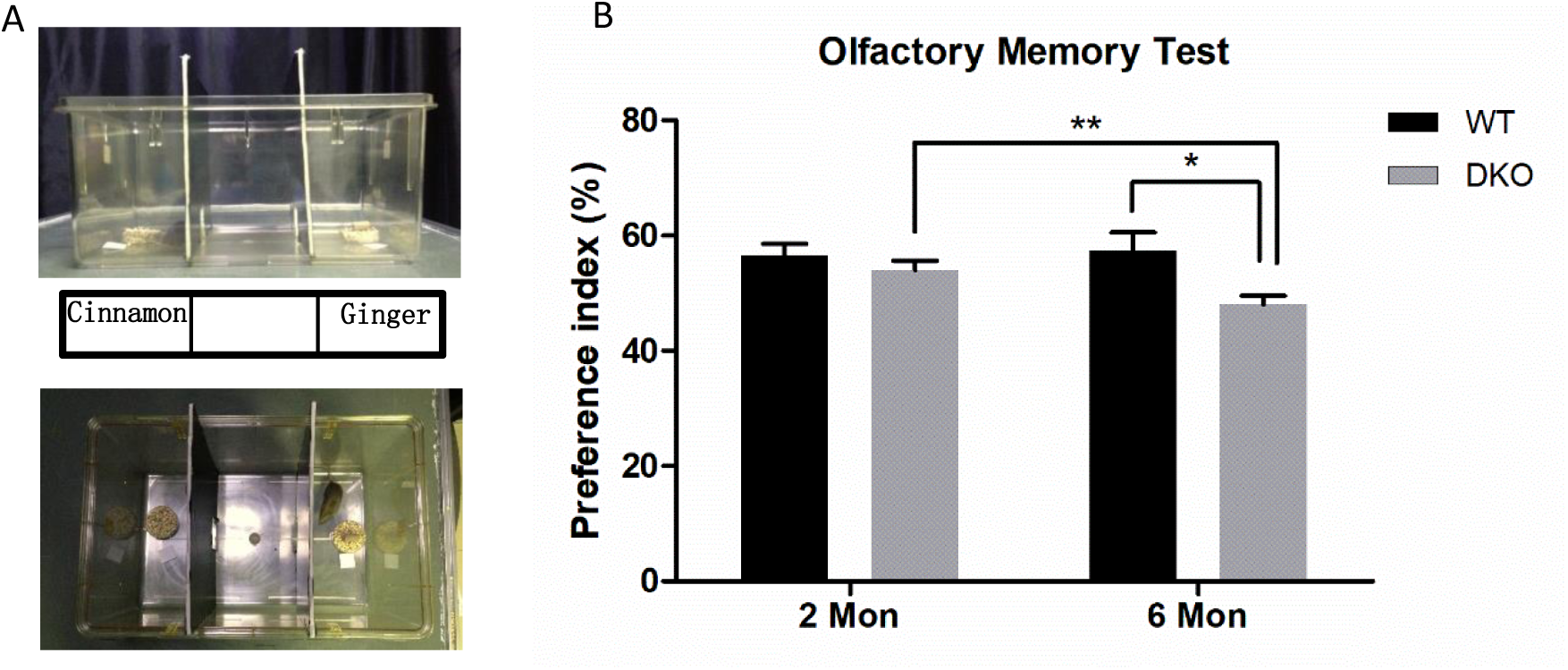
Olfactory memory test in 2-month-old and 6-month-old WT and DKO mice. A. The apparatus. The filter paper dipped with cinnamon was placed in the left compartment, and the filter paper dipped with ginger essential oil was placed in the right compartment together with the food pallet at the training phase. At the test phase, the food pallet was removed. B. Olfactory memory test results. The percentage of the time mice spent in the right compartment was compared. ***, P < 0.001.

**Figure 4.**
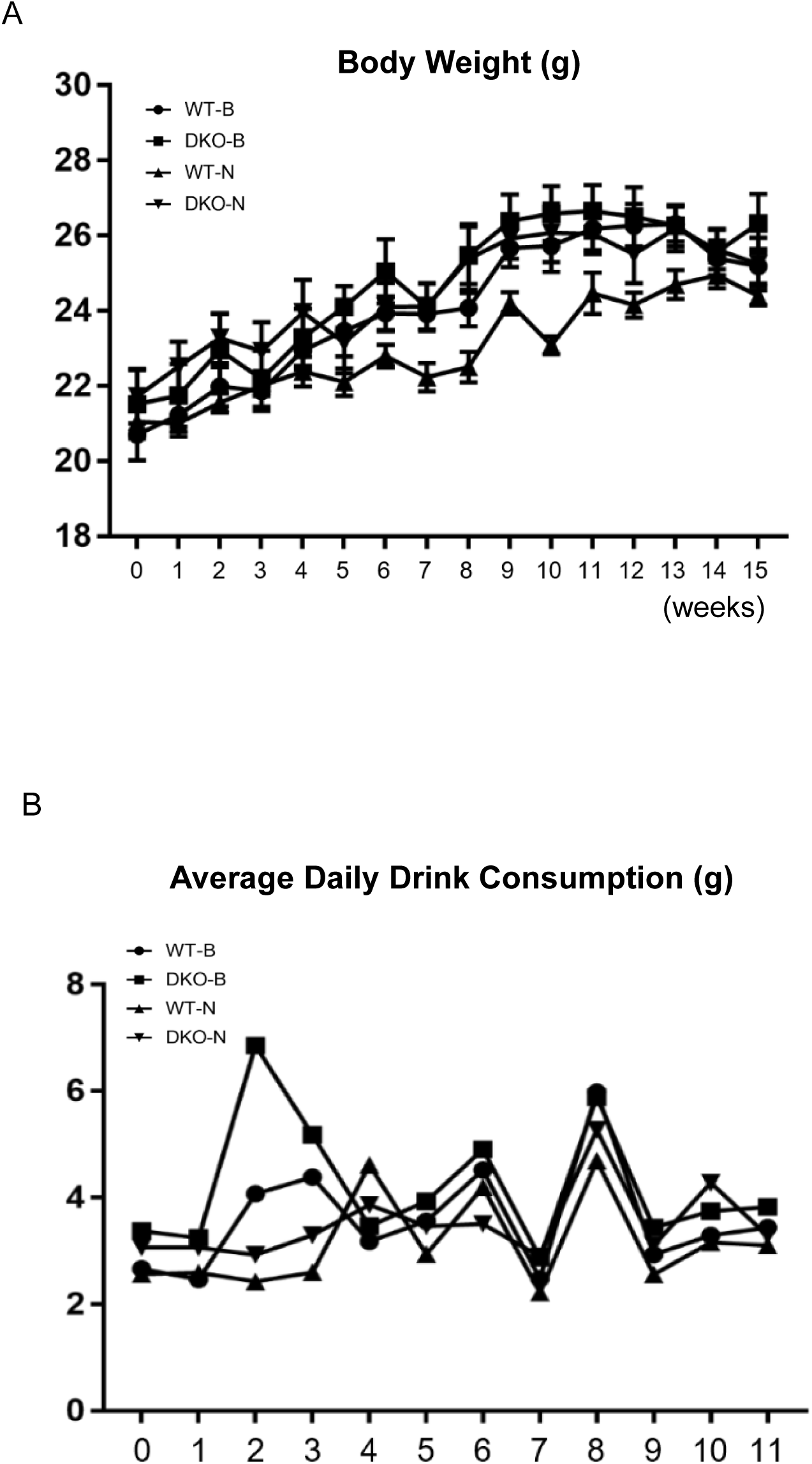
Mice body weight and drinking consummation measurement during the nicotine administration in WT-B, DKO-B, WT-N and DKO-N group of mice. A. Body weight curve. B. water drinking consumption curve.

After a 4-month nicotine administration period, mice underwent handling for three consecutive days, with 5 minutes per day per mouse, to acclimate them to the experimenters. Olfactory functional behavior tests were then conducted to assess the impact of nicotine on both WT and DKO mice. Statistical analysis indicated no treatment effect in any of the three tests: the nicotine effect *F* _(1.39)_ *=* 0.131, P = 0.720 in the buried food test; *F* _(1.39)_ *=* 0.137, P = 0.714 in the olfactory avoidance test, and *F* _(1.39)_ *=* 0.218, P = 0.643 in the olfactory memory test. The genotype effect was observed in the buried food test (*F* _(1.39)_ *=* 9.716, P = 0.004) and the olfactory avoidance test (*F* _(1.39)_ *=* 4.534, P = 0.04).

In detail, in the buried food test, DKO-B mice spent significantly more time retrieving the food pellet than WT-B mice (83.2 ± 12.3 sec Vs 215.3 ± 66.3 sec, P < 0.05); after nicotine administration, the DKO-N mice also showed prolonged latency compared to WT-N mice (243.3 ± 74.6 sec Vs 88.5 ± 14.6 sec, P < 0.05) (Figure 5 A). In the olfactory avoidance test, WT-B mice displayed an avoidance time of 30.3 ± 6.5 sec, while DKO-B mice showed almost no avoidance (0.4 ± 8.1 sec, P < 0.01), suggesting the olfactory memory deficiency in DKO mice at 6-month-old. Nicotine-treated groups showed no significant differences, with WT-N mice avoiding the odor for 22.0 ± 9.0 sec and DKO-N mice for 15.1 ± 11.5 sec (P > 0.05) (Figure 5B). In the olfactory memory test, the preference index of WT-B mice was 66.7 ± 4.1%, while DKO-B mice showed a significantly lower index of 51.7 ± 5.4% (P < 0.05), indicating the olfactory memory deficiency in DKO mice at 6-month-old. With nicotine treatment, the preference indices were 52.0 ±11.2% in WT-N mice and 59.9 ± 3.8 % for DKO-N mice(P > 0.05), showing no improvement. (Figure 5 C). When comparing WT and DKO mice with nicotine treatment (WT-N and DKO-N), no significant differences between these two groups of mice in either the olfactory avoidance or olfactory memory tests, suggesting that nicotine administration did not improve olfactory function in DKO mice.

**Figure 5.**
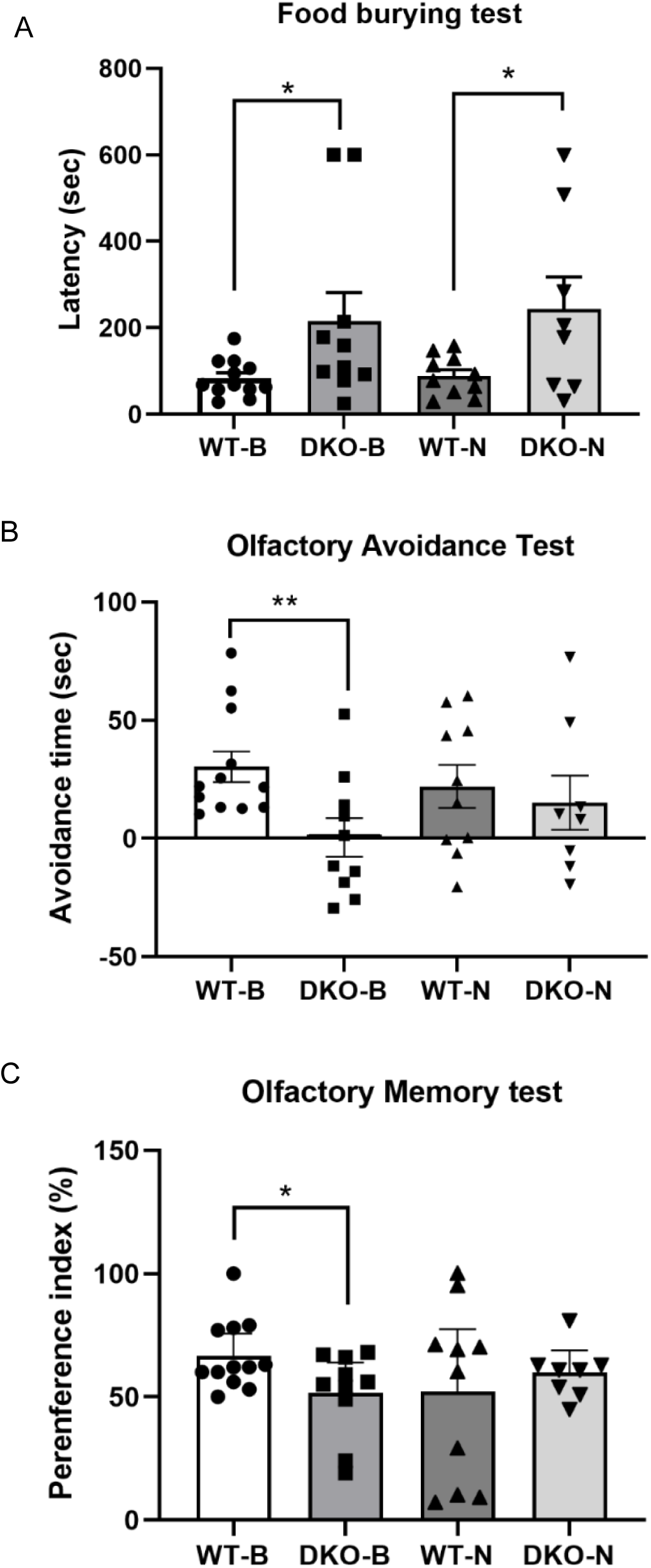
Nicotine effect on olfactory functional behaviors in WT and DKO mice after 4-month nicotine administration. A. Buried food test. DKO mice showed deficiency in digging the buried food pallet compared to WT mice (p < 0.05), with or without nicotine administration. B. Olfactory avoidance test. DKO-B mice showed declined odor recognition capability compared to WT mice. After nicotine administration, there was no significant difference between WT-N and DKO-N mice; there was also no significant difference when comparing DKO-B to DKO-N mice or comparing WT-B to WT-N mice. C. Olfactory memory test. DKO-B mice showed deficiency of olfactory memory capability compared to WT-B mice. After nicotine administration, there was no significant difference between WT-N and DKO-N mice; there was also no significant difference when comparing DKO-B to DKO-N mice or comparing WT-B to WT-N mice.

It is notable that although no significant differences were observed between WT-B and WT-N mice, a trend of declining olfactory function was seen in WT-N mice. Comparison between WT-N and DKO-B mice revealed no significant differences (P = 0.08 in the buried food test; P = 0.09 in the olfactory avoidance test; P = 0.97 in the olfactory memory test), suggesting that chronic nicotine administration may have a potential negative effect on olfactory function even in healthy young mice.

## 4. Discussion

Olfactory dysfunction is a prevalent pathological feature in neurodegenerative diseases, albeit with varying degrees of impairment due to differences in underlying neural damage. The olfactory bulb receives input from the central cholinergic system, and cholinergic axons from the basal forebrain modulate olfactory synaptic activity. Consequently, the correlation between performance in olfactory tasks and forebrain cholinergic denervation has been extensively explored in a broad spectrum of neurodegenerative diseases, with particular emphasis on AD. Wilson et al. found that the density of neurofibrillary tangles was the main pathological factor that affected olfactory function, especially in the entorhinal cortex, hippocampus CA1 region, and subiculum; after controlling for tangles, the effect of amyloid burden was attenuated ^26^. In APP mutant overexpression 3×Tg-AD mice, researchers correlated Aβ deposition in olfactory bulb with olfactory dysfunction ^27^. DKO mice develop Tau hyperphosphorylation but without Aβ deposition which supports the non-amyloidogenic mechanism and provides a unique model for further study of the mechanical relationship between olfactory function and neurodegeneration.

Clinical studies show early olfactory deficits, including impaired detection and identification thresholds, in individuals with AD before full clinical onset ^28^. These findings align with our results, where DKO mice exhibit deficiencies in odor detection capabilities at 2 months of age, despite their odor discrimination and olfactory memory remain normal. Furthermore, cognitive and non-cognitive species-specific behaviors show no significant changes, further suggesting that olfactory assessment could serve as an early diagnostic method for detecting AD. Owing to its convenience and cost-effectiveness, an olfactory detection assay could prove to be a valuable diagnostic tool. The progression of neurodegeneration in DKO mice, reflected in worsening detection, discrimination, and memory deficits by 6 months, emphasizes olfactory dysfunction as an early marker of disease ^29^. Interestingly, no sex differences were observed in olfactory behavior, despite clinical evidence suggesting women are more susceptible to AD.

The function of nicotine, a nicotinic acetylcholine receptor agonist, in the regulation of olfactory function and/or neurodegenerative diseases is complex and contested. Nicotine can enhance the odor-induced increase in response of olfactory bulb blood flow (but not the basal blood flow level) ^30^. In vivo chronic nicotine exposure significantly decreased the number of newborn granule cells in adult wildtype mice, but since an increase in the number of granule cells does not necessarily correlate with better olfactory performance, it is still unclear whether the chronic nicotine exposure can enhance or inhibit the olfactory function in healthy adult mice ^31^. Conflicting results from amyloidogenic transgenic mice have shown significant Aβ-lowering effects, or alternatively, no effects of nicotine administration ^32-34^. Our results showed that chronic nicotine exposure failed to rescue the neurodegeneration in young adult DKO mice and might suggest a potentially negative effect in young WT mice. When summarizing these studies (see Table 1), we noticed that age might be an inevitable factor. Connor et al. compared the nicotine effect in adolescence and adulthood and found that after over 12 days of nicotine administration, short-term (24 h) or long-term (30 days) abstinence elicited the opposite effect on learning and memory behaviors in adolescents and adults ^35^.

**Table 1.** Nicotine effect in AD animal models.

| AD model | Starting Age | Nicotine administration | Effect | Reference |
| --- | --- | --- | --- | --- |
| Tg2576 (APP<sup>sw</sup>) mice | 9-month-old | 10-days | Aβ↓ | <sup>37</sup> |
| Tg2576 (APP<sup>sw</sup>) mice | 9-month-old | drinking fluid (200 mg/ml) for 5.5 months | Insoluble Aβ1-40 and Aβ1-42↓;<br>soluble 1-40 or 1-42 unchanged | <sup>38</sup> |
| Tg2576 (APP<sup>sw</sup>) mice | 10-month-old | Gradually from 0.25 mg/kg to 0.45 mg/kg (i.p), totally 10 days | guanidinium-soluble Aβ↓.<br>intracellular Aβ unchanged | <sup>39</sup> |
| 3×Tg-AD mice | 1-month-old | Gradually increased profile: 25-600 μg/ml, 5 months in total | aggregation and phosphorylation state of tau↑ | <sup>32</sup> |
| 3×Tg-AD mice | 3-month-old | 200 μg/ml via drinking for 20 weeks | No change of Aβ level | <sup>40</sup> |
| Streptozotocin AD model rat | 35 - 42 days | 1 mg/kg (i.p) for 20 days | Attenuated behavioral and synaptic plasticity impairments | <sup>34</sup> |
| Fisher 344 rats | 4-month - old and 24-month-old | 0.2 mg/kg (i.p), 15min before behavior tests | Improve working memory in young rat, no beneficial effect in aged rat | <sup>41</sup> |
| NMRI mice | 2, 6, 10-month-old | 0.35 mg/kg for 10 days | No improvement in Water maze | <sup>42</sup> |

Interestingly, nicotine’s effect on PD seems more beneficial. For example, one-week preventive and the following 2-weeks (after MPTP injection) nicotine administration was reported to improve the olfactory impairment in a mouse model of Parkinson’s disease ^33^. However, in cases of ischemic injury, nicotine exacerbated brain damage ^36^, further illustrating the complexity of nicotine’s effects, which may depend on factors like age, health, and disease state.

In summary, our results demonstrated that Presenilin-1/2 double knockout impairs odor detection in young adult mice and accelerates the progression of olfactory discrimination and memory impairment. The mechanism of Presenilin-1/2 in regulating olfactory function warrants further investigation, as early detection deficits appear to correlate with neurodegeneration. Moreover, chronic nicotine administration was not found to have a preventive effect on mitigation olfactory function deficiency in young adult DKO mice; however, nicotine administration in DKO with clinical symptoms is still worthy of exploration in the future.

While our data indicate that mice demonstrate a clear preference to avoid areas containing 2-mercaptobenzothiazole (2-MB) compared to water, suggesting an odor-driven behavior, an essential consideration highlights: the longer time spent in the larger compartment during habituation might suggest a complexity that extends beyond simple olfactory aversion. This behavior could be influenced by a variety of factors, including a preference for more spacious environments or innate exploratory behaviors, underscoring a potential limitation in our methodology that deserves acknowledgment.

Our findings primarily demonstrate an ability to detect and react to aversive odors, which, while indicative of olfactory sensitivity, do not necessarily speak to the animals’ capacity for odor recognition or discrimination. This distinction is crucial for interpreting our results within the broader context of olfactory function research. Therefore, it is important for us to clarify that while our study provides insight into the olfactory capabilities of mice, particularly their aversion to certain odors, it may not fully capture the nuances of olfactory perception, including memory and differentiation of odors. Future studies are encouraged to employ additional behavioral assays that can more directly assess these complex aspects of olfactory function.

## Ethics in publishing

All experiments were approved by the Institutional Animal Care and Use Committee of the East China Normal University (IACUC approval ID #M10020).

## Acknowledgement

This work was financially supported by the grant from the Joint Institute of Tobacco and Health (No.2022539200340036).

## Declare of Interests

It is important to note that the financial sponsor had no role in the study design, data collection and analysis, decision to publish, or preparation of the manuscript. The decision to publish and the content of the manuscript are solely the responsibility of the authors.

Feiyan Qi is an employee of the Joint Institute of Tobacco and Health. Her contribution to this study was limited to providing general advice on research direction. She was not involved in the design, execution, data analysis, or interpretation of the experiments, nor did she influence the preparation of the manuscript. The authors declare that there are no conflicts of interest that influenced the outcomes or conclusions of this study.

All other authors have no conflict of interest to report.

